# Real-time Sub-cycle Oscillatory Beta Burst Detection

**DOI:** 10.64898/2026.05.19.726185

**Authors:** Anirudh Wodeyar, Joël Karel, Ralf Peeters

## Abstract

Beta-band burst activity is a key biomarker of Parkinsonian pathophysiology and a control signal for adaptive deep brain stimulation (aDBS). Existing burst detection approaches rely on bandpass filtering followed by envelope extraction and thresholding, which introduces onset/offset latency due to sliding-window estimation and can be unstable under variations in signal-to-noise ratio or threshold choice. We introduce a switching state space approach for real-time burst detection that models oscillatory activity as a superposition of latent harmonic components and explicitly represents burst and non-burst regimes. The framework performs real-time inference using a set of Kalman filters and computes posterior mode probabilities using a Markov transition prior, producing sample-by-sample burst probabilities together with phase and uncertainty estimates. In simulated data with sinusoidal beta bursts embedded in both white and pink noise, the method improved burst detection accuracy, reduced onset latency and offset latency compared with causal envelope-based baselines, and showed reduced sensitivity to threshold selection. In analysis of real sub-thalamic nucleus recordings during a grip-force task, the MSSR method recovered movement-related burst modulation consistent with prior reports. Our results indicate that state space switching provides a principled route to low-latency burst detection that may better support closed-loop stimulation strategies timed to beta bursts.

## I. Introduction

Parkinson’s disease (PD) is frequently characterized as an oscillopathy, with prominent alterations in beta-band activity (13–30 Hz) in cortico-basal ganglia circuits [1]. Beta-band activity tends to occur transiently, with amplitude waxing and waning on short timescales [2]. In recordings from the subthalamic nucleus (STN), beta-band power correlates with motor impairment and changes systematically with clinical state [3]. Pharmacological treatment with levodopa commonly reduces the beta peak in the STN power spectrum in responsive patients, and continuous deep brain stimulation (DBS) similarly suppresses beta-band activity [3]. Together, these findings have supported the hypothesis that beta dynamics are not merely epiphenomenal, but may serve as a clinically relevant control signal for therapy.

This motivation has contributed to the development of beta-timed adaptive DBS (aDBS) strategies in which stimulation is delivered contingent on beta-band activity [4]–[6]. Such systems can reduce total stimulation energy and may mitigate stimulation-related side effects. In many implementations, online beta detection is performed using bandpass filtering, envelope extraction, and thresholding. While effective in offline analyses, these approaches face two practical limitations for real-time (streaming) burst-triggered control. First, windowed filtering (between 0.1 and 0.4 s long) and envelope estimation introduce unavoidable latency, which can delay both detection of burst onset and offset and thus, cause stimulation to persist beyond burst termination. Beta bursts can sometimes be as short as 0.1 s, which can be substantially attenuated or missed due to smoothing and delay. Second, performance depends strongly on the choice of detection threshold, which can vary across subjects, recording conditions, and signal-to-noise regimes. These issues are particularly problematic when stimulation timing is intended to align closely with the true onset and offset of beta bursts.

To address these limitations, we build on our prior work [7] demonstrating that rhythmic activity can be tracked in real-time using a state space model of complex-valued damped harmonic oscillators driven by stochastic inputs. This modeling framework provides a structured prior for oscillatory dynamics and, because it is linear-Gaussian, supports efficient sample-by-sample inference via the Kalman filter. Here, we extend this approach by introducing a switching state space model inspired by hidden Markov models and multiple-model adaptive estimation [8]. We treat bursts as latent regime changes and run two competing linear dynamical models in parallel: one containing a beta-band oscillator component and one without. By performing real-time inference and updating mode probabilities at each sample, the proposed method yields a continuous-valued estimate of burst probability, alongside phase and uncertainty estimates [9], enabling low-latency and threshold-robust detection suitable for closed-loop neuromodulation.

## II. Methods

### A. State Space Model of Rhythms (SSR)

We modeled the observed neural time series *y*_*t*_ as a sum of *K* rhythms plus additive measurement noise. Each rhythm *j* is represented by a two-dimensional latent state 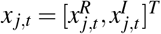, corresponding to the real and imaginary components of the analytic signal.

State evolution (stacking all rhythms into *X*_*t*_ ∈ ℝ^2*K*×1^):

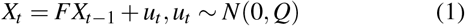

where *F* is block-diagonal with 2× 2 blocks *F*_*j*_ = *a* _*j*_*O*(*ω*_*j*_) and *Q* is 2*K*× 2*K* matrix for the noise covariance. Here *O*(*ω*_*j*_) is a rotation matrix with angular frequency *ω*_*j*_ = 2*π f* _*j*_*/F*_*s*_, and 0 *< a* _*j*_ *<* 1 controls damping/stability.

Observation model:

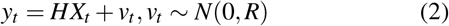

where *H* = [1, 0, 1, 0, …, 1, 0] selects and sums over the real components of all rhythms, *R* is a scalar representing the measurement noise variance. Given (1)–(2) and Gaussian noise assumptions, we used a Kalman filter to obtain the posterior state estimate 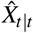 and covariance 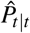 for each incoming sample. Instantaneous amplitude and phase for rhythm j were computed as:

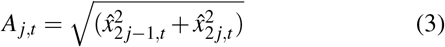

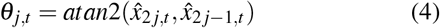

### B. Multiple State Space Model of Rhythms (MSSR)

To detect beta bursts online, we formulated a two-mode multiple-model adaptive estimation problem (see Fig. 1 for the architecture). Mode *s*_*t*_∈ {0, 1} indicates absence/presence of a beta-band oscillator. We constructed two SSR models: *M*0 excluded the beta oscillator and *M*1 included it. Both models included all non-beta oscillators and were propagated in parallel through independent Kalman filters.

**Fig. 1.**
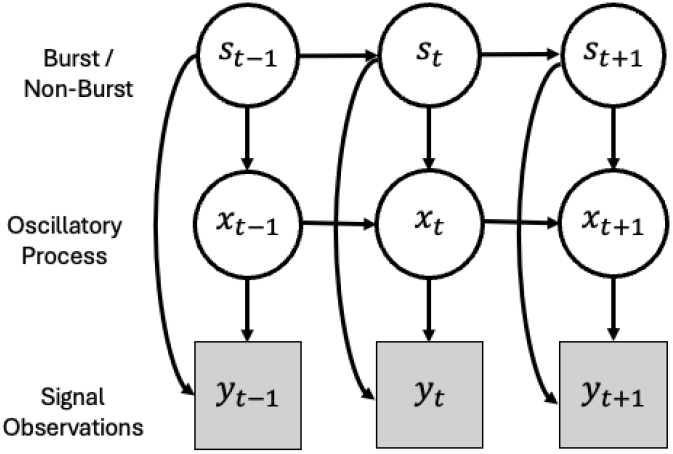
Model Architecture. The occurrence of bursts is tracked as a latent state *s*_*t*_ which modulates both the state (*x*_*t*_) equations as well as the expected observations (*y*_*t*_). Both the probability of the burst *s*_*t*_ and the oscillatory state *x*_*t*_ are propagated through time according to distinct equations.

Mode probabilities *µ*_*t*_ = [*P*(*s*_*t*_ = 0), *P*(*s*_*t*_ = 1)]^*T*^ evolve according to a first-order Markov chain with transition matrix *T*. The predicted mode distribution is given as:

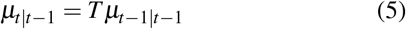

For each model, we computed an innovation likelihood *L*_*t*_ based on its predicted measurement distribution:

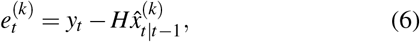

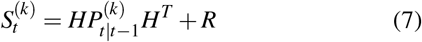

and updated the posterior mode probabilities via Bayes’ rule:

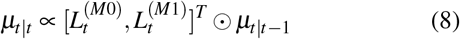

followed by normalization so that mode probabilities sum to one. The beta-burst probability was defined as *p*_*β*_ (*t*) = *P*(*s*_*t*_ = 1 | *y*_1:*t*_). We did not mix state estimates across models as we found that state mixing degraded tracking performance due to the oscillator-based state representation.

Although the switching state-space formulation is not inherently restricted to online use, the present work focuses on its causal filtering implementation. In this implementation, inference at time *t* depends only on observations available up to that sample.

### C. Parameter initialization

We estimated SSR parameters (frequencies *f*_*j*_, damping *a*_*j*_, and noise covariances *Q* and *R*) acausally from an initial 10*s* segment using an EM-based fitting routine (MATLAB implementation). The beta oscillator was identified as the fitted rhythm with frequency within 13*−* 30 *Hz*.

To initialize the Markov transition matrix *T*, we computed the beta amplitude envelope 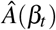 from the acausal beta state estimate using (3) and discretized it into quartiles. We then fit a two-state hidden Markov model (burst/non-burst) with four emission bins using Baum–Welch (MATLAB hmmtrain), with initial emission probabilities that placed more mass on high-amplitude bins for the burst state and on low-amplitude bins for the non-burst state (specifically [1/3, 1/3, 1/4, 1/12] for non-burst and [1/12, 1/12, 1/3, 1/2] for burst across the four amplitude quartile bins).. This effectively enabled more emissions of high amplitude instants in the burst state.

During online filtering, we inflated the observation noise co-variance (*R →* 2*R*) to increase robustness to model mismatch.

### D. Contemporary Methods Implementation

#### 1) Causal FIR

As comparison method to the MSSR approach, we implemented a real-time method that parallels existing contemporary algorithms [4]. We apply a 0.3 s sliding window updated each sample (most recent sample at the window’s end) and zero-phase filter the window contents using an FIR filter of order 77 (to reasonably capture power at the lowest potential frequency of 13 Hz). We use a sliding window ending at the current sample and report the envelope estimate aligned to the window endpoint. Although filtering within the window is acausal, the overall pipeline produces a streaming estimate without access to future samples beyond the current window endpoint. We apply the Hilbert transform to this filtered signal and generate the analytic signal (using MATLAB’s *hilbert* implementation). From the analytic signal, we calculate the absolute value and take the average over the final 0.1 s as an estimate of the beta-band amplitude for the most recent sample of data. This helps partially correct for the edge error from the Hilbert Transform application. This amplitude estimate is converted to a percentile and used to ascertain presence of a burst.

#### 2) Forecast FIR

To account for the inherent latency of the Causal FIR approach, we modified an existing method [10] for real-time phase estimation to enable burst estimation. This method first estimates the parameters of an autoregressive (AR) model for each window of data, then forecasts 0.1 s of data, before finally applying a filter and then the Hilbert transform to estimate amplitude and phase. We modify the version of the Zrenner et al. (2020) [10] algorithm accessible at https://github.com/bnplab/phastimate. Please refer to Zrenner et al. (2020) for algorithm details. Compared to the default settings, we set the algorithm to track a beta band rhythm by changing the window size for the filter to 0.3 s, the filter order to 77, and the frequency band to 13 to 30 Hz. Finally, we average the final 0.2 s of the window (0.1 s on either side of the most recent sample of data) to get an estimate of the current amplitude. Amplitude is then converted into percentiles. A threshold is applied to determine whether a burst is occurring.

### E. Simulations

To test the methods, we implemented a novel simulation that enabled access to ground truth while reflecting the statistical structure of data. We set up bursts of beta band oscillations to be sinusoids that vary from burst to burst in amplitude, starting phase and center frequency. The amplitude range was constrained between [2, 15], starting phase was uniformly random over [0, 2*π*) and center frequency *C*_*f*_ *∼N* (16 Hz,1 Hz). Burst durations were modeled with a truncated Exponential distribution with mean 0.3 s, with minimum burst duration set to 0.05 s and maximum duration to 1 s. This is in accordance with expected durations for beta bursts [3]. Time between bursts was similarly modeled with a truncated Exponential distribution with a mean of 0.4 s and with minimum refractory period of 0.25 s. Note that this approach mimics a point-process with refractory periods for the beta burst process. To this base signal we added either white or pink noise (spectral density follows 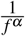 for frequencies *f* and *α* = 1) whose standard deviation was set to 4 as the measurement noise. An example trace of the sinusoidal bursts in pink noise is shown in Figure 2. All simulations sampled data at 1000 Hz to replicate expected sampling rates in clinical settings.

**Fig. 2.**
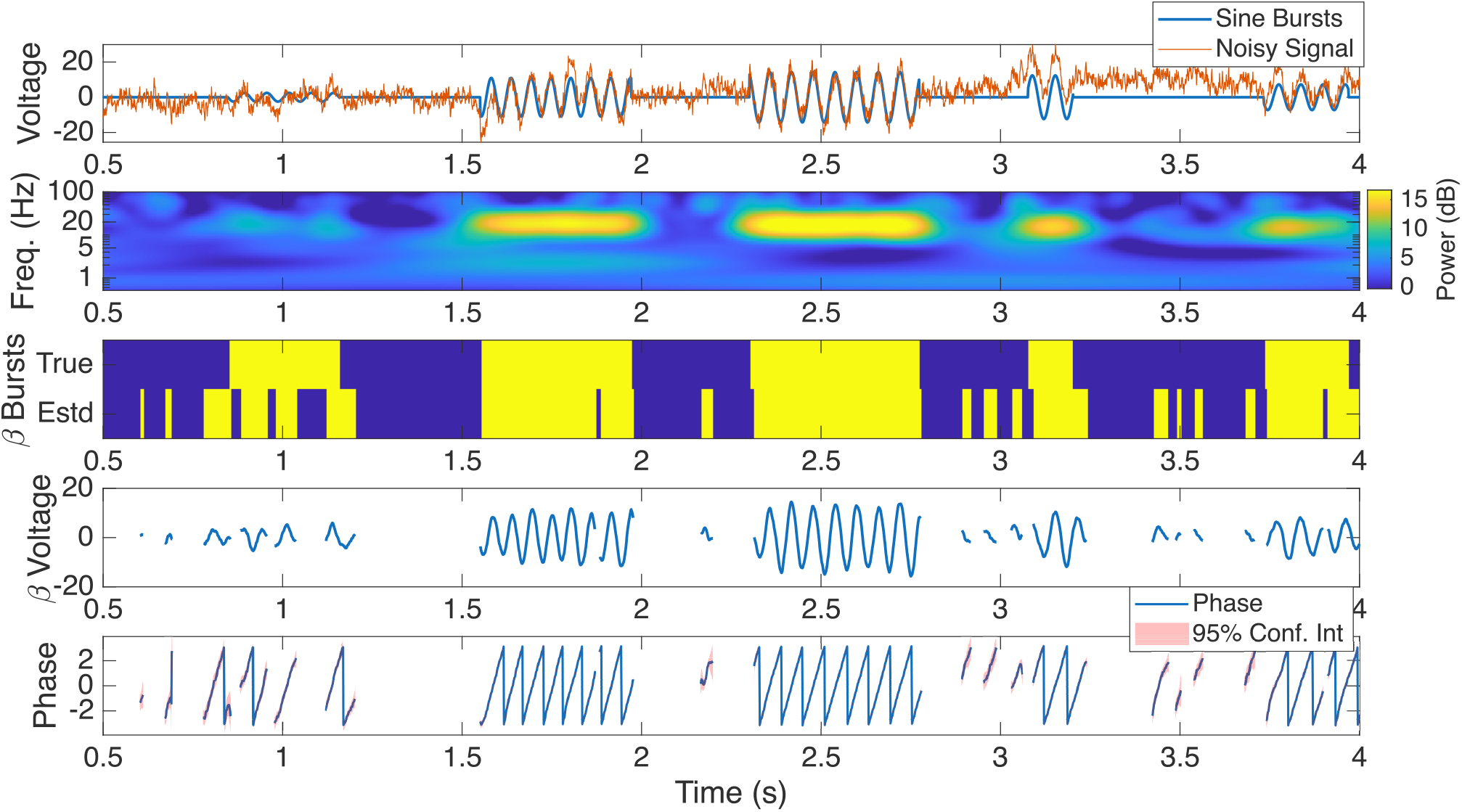
Example of beta burst detection using MSSR. Top row shows the pure sine bursts (blue) and the noise corrupted signal (orange). Second row shows the smoothed wavelet spectrogram of the noise-corrupted signal. Third row shows periods where bursts are present (True) and the estimated periods of the bursts using the MSSR model (Estd.). Note that even the very low amplitude initial burst is reasonably captured and latency for detecting middle two higher amplitude bursts is near zero. Bottom two rows demonstrate capability of MSSR to extract both the burst activity and the phase simultaneously, and to provide uncertainty estimates.

### F. Metrics

We measured F1 score, and latency at detection and offset. F1 score helps balance precision and recall, critical for the application scenario we target here. Precision is the ratio of true positives to the total set of positive predictions. A true positive is a sample that contains a burst that is also labeled by an algorithm to contain a burst. Recall is the ratio of true positives to all positives that existed in the ground truth. To reduce noise in latency assessment, we dropped periods deemed bursts that were shorter than 17 ms as this corresponds to half a cycle at 30 Hz which is too short to expect a true burst.

### G. Data

We evaluated MSSR on an openly available intraoperative dataset from patients with Parkinson’s disease undergoing STN-DBS implantation [11]. The original dataset contains simultaneous subthalamic local field potential recordings and sensorimotor electrocorticography recorded while patients performed a cued Go/No-Go grip-force task. Experimental procedures involving human subjects were approved by the Institutional Review Board of University of Pittsburgh. In the present proof-of-concept analysis, we used only the STN recordings.

For one representative subject, STN contacts were analyzed in a bipolar montage, and the bipolar channel with the clearest spectral peak in the beta range was selected based on visual inspection. Model parameters were initialized from an initial segment of data, after which burst probability was estimated sample by sample as the data was continuously streamed to the model.

Movement onset times were obtained from the task/force annotations and were used only for post hoc alignment of the estimated burst probability. Burst probabilities were averaged across valid movement trials for visualization of movement-related beta-burst modulation.

## III. Results

We first evaluated MSSR on the pink-noise simulation (Fig. 2). SSR parameters were estimated acausally from the first 10 s of each trace, and the Baum–Welch algorithm was then used to estimate the two-state transition matrix from the SSR-derived beta-band amplitude. The resulting switching model was subsequently applied online (data was streamed to the models) to a held-out 10 s segment. In the representative example shown in Fig. 2, MSSR tracked burst onset and offset with minimal delay once the burst amplitude exceeded the noise standard deviation (*σ* = 4). In addition to burst probability, the model provides instantaneous phase estimates and posterior uncertainty bounds, which can be used to restrict stimulation to time points with high phase certainty.

We next compared MSSR with the Causal FIR and Forecast FIR baselines on the same simulated trace (see example in Fig. 3). For each method, as an initial test, we selected the decision threshold that maximized the sample-wise F1 score for that trace (oracle threshold). The oracle threshold was established on the probabilities output from the MSSR and the amplitude percentiles from the Causal FIR and Forecast FIR. Later, we report results from examining performance over a range of thresholds. Given the oracle threshold, MSSR detected low-amplitude burst structure and, critically, identified the onset of the two high-amplitude bursts without the systematic delay observed for the FIR-based approaches. As expected, the Forecast FIR method reduced onset latency relative to Causal FIR, but both retained unavoidable delays consistent with their use of sliding-window envelope estimation. Qualitatively, we observed that the starting phase of a burst influenced detection latency for both FIR baselines, whereas MSSR performance was insensitive to starting phase.

**Fig. 3.**
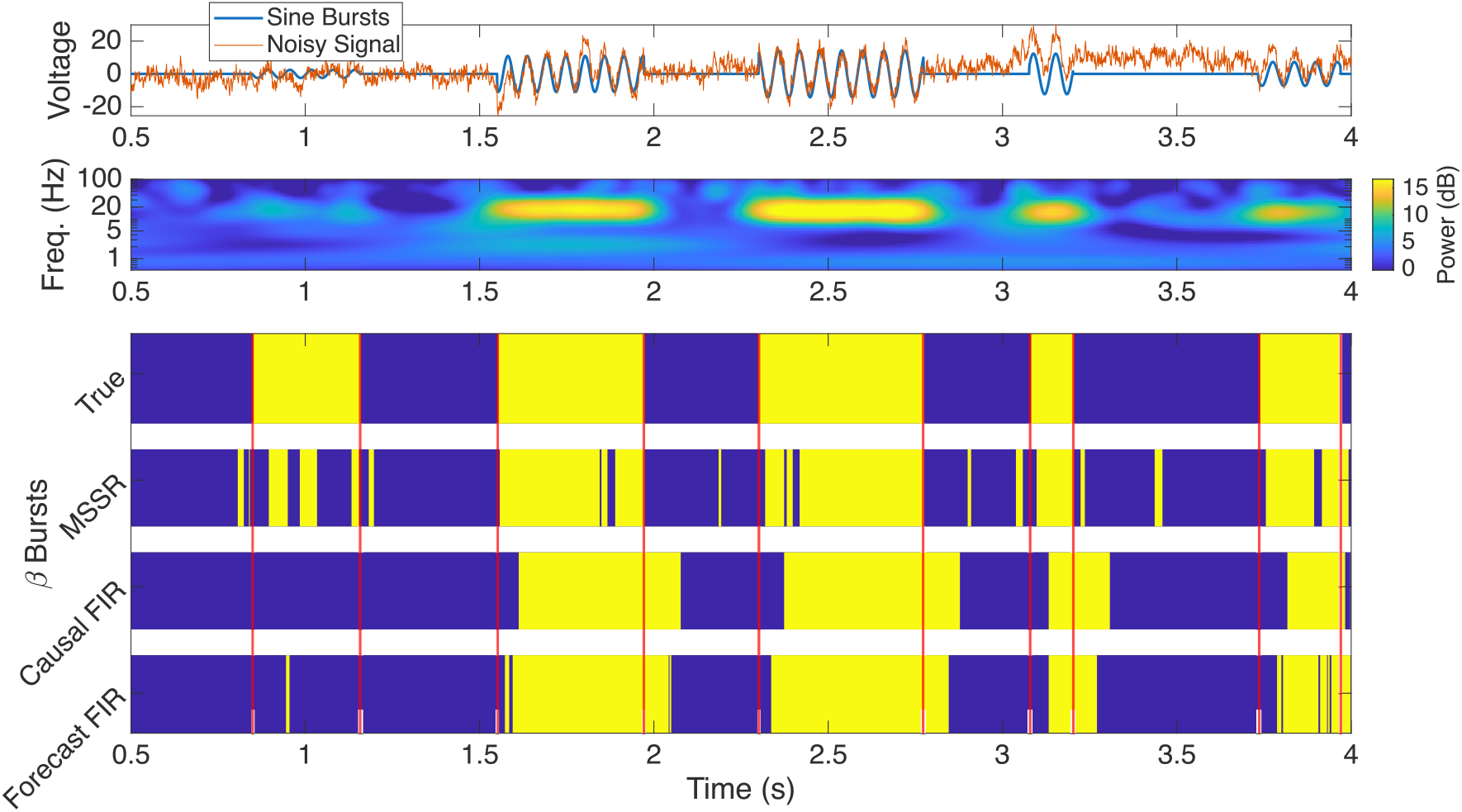
Comparison of methods when data is streamed. Top row shows signal being tracked, with the time-frequency spectrogram immediately below. Lower set of subplots shows moments of burst being detected (yellow) and period without bursts being detected (blue) for three methods applied to streamed data.

To quantify performance across realizations, we repeated each simulation condition (bursts in white noise and bursts in pink noise) for 200 trials. For each trial and method, we computed the sample-wise F1 score and burst onset/offset latencies relative to ground-truth burst boundaries, using a minimum predicted burst duration of 17 ms to suppress spurious detections and only including bursts where at least 50% coverage was achieved. Note that, methods marking the entire signal as a burst have a baseline F1-score of 0.6. In white noise, MSSR achieved a mean oracle F1 score of 0.93 (SE 0.002), with onset latency 22.53 ms (SE 0.56) and offset latency 14.6 ms (SE 0.33). Both FIR baselines showed significantly lower oracle F1 scores (0.74 for Causal FIR and 0.81 for Forecast FIR; paired t-tests, p < 0.05) and larger onset delays (49.67 ms and 36.28 ms, respectively; two-sample t-tests with unequal variances, p < 0.05) and larger offset delays (121.85 ms and 75.31 ms, respectively; two-sample t-tests with unequal variances, p < 0.05).

When methods were evaluated on pink noise, all methods degraded but the same trends were preserved (Fig. 4). For the oracle threshold, MSSR reached a mean F1 score of 0.86 (SE 0.003), onset latency 13.12 ms (SE 0.37), and offset latency -12.06 ms (SE 0.46). A negative offset latency indicates that the method deemed the burst to end earlier than the true burst ending. MSSR had an offset latency up to 100 ms faster than the baselines. Both FIR baselines showed lower oracle F1 scores (Causal FIR: 0.72 and Forecast FIR: 0.8, p < 0.05), larger onset delays (Causal FIR: 54.08 and Forecast FIR: 32.51; p < 0.05) and larger offset delays (Causal FIR: 114.77 and Forecast FIR: 75.02; p < 0.05). Critically, MSSR remained robust across a broad range of decision thresholds (F1 > 0.7), whereas both FIR baselines were highly sensitive to threshold selection: their F1 curves peaked sharply near the oracle percentile threshold and dropped rapidly for nearby values (Fig. 4, top right). Thus, in simulations, the method improved detection accuracy while reducing onset and offset latency relative to causal envelope-based baselines. However, a potential limitation is that the comparison was restricted to a causal envelope-based baseline (albeit, one that reflects contemporary practice) and one forecasting variant; future work will compare MSSR against a broader set of optimized burst-detection methods.

**Fig. 4.**
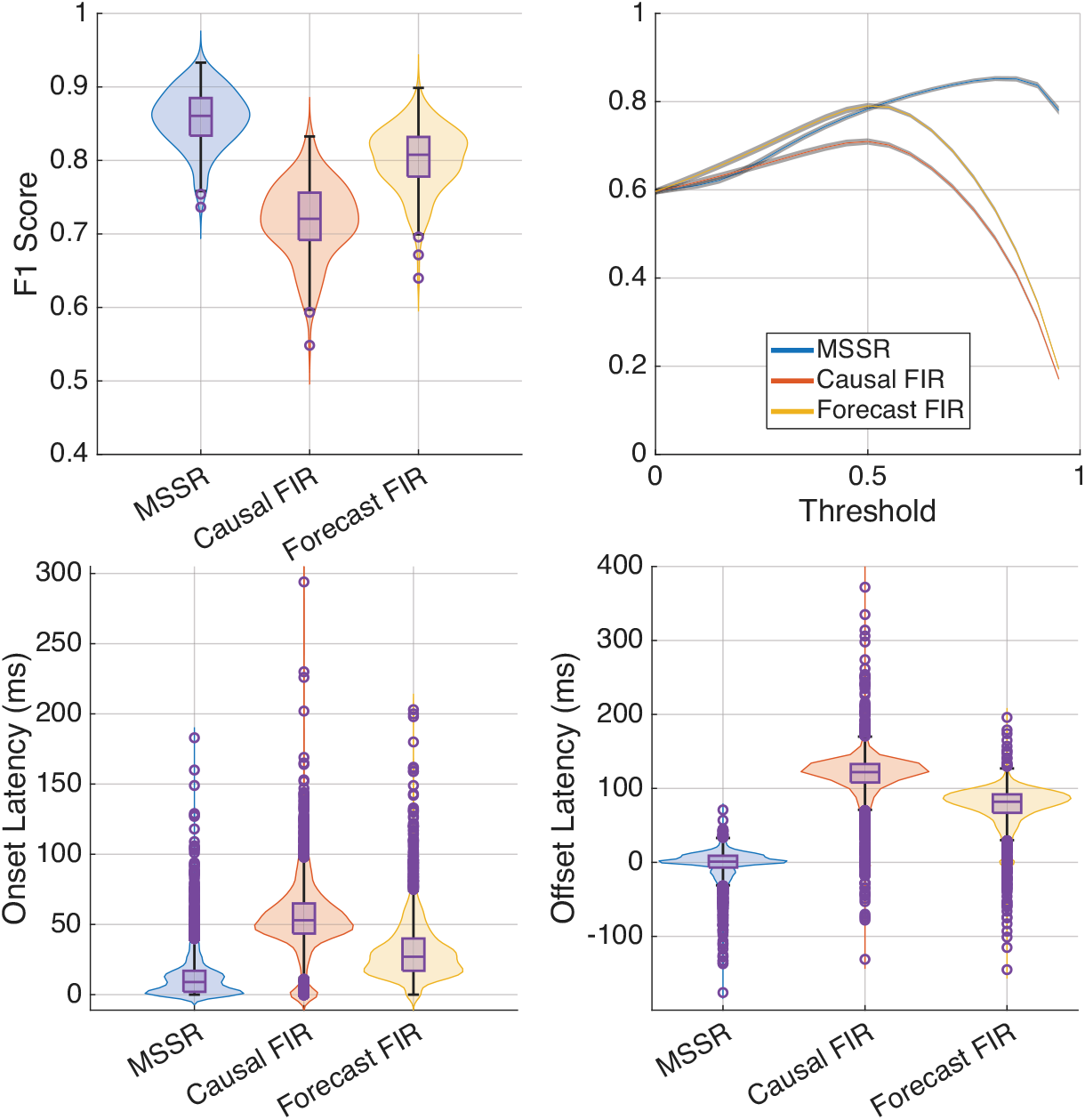
MSSR has higher F1 scores and lower latencies than baseline methods for sinusoidal beta bursts in pink noise. The top row shows the F1 score and bottom row shows the latencies of onset and offset detection. Top left shows the oracle F1 score (each dot is one simulation) demonstrating consistently better performance of the MSSR. Top right shows F1 score with 95% CI across a range of thresholds (either probability for MSSR, or amplitude, for causal FIR and Forecast FIR). Bottom row each dot represents a burst, with the distributions modeling latencies across all bursts. Note that we have cut the y-axis off (some estimates of the Causal FIR are not visible) to enable easier comparisons.

Finally, we evaluated MSSR on real neural recordings using an openly available dataset [11]. For one subject, we analyzed subthalamic nucleus (STN) local field potentials time-locked to hand-movement onset, replicating a previously reported movement-related beta modulation. Based on qualitative inspection of the power spectral density, we modeled the signal using two oscillatory components: one in the delta range (0.5–2 Hz) and one in the beta range (13–30 Hz). The remainder of the processing pipeline matched the simulation experiments. MSSR was computationally efficient in this setting, requiring 0.0001 s per Kalman prediction–correction update on an Apple MacBook M3 Pro, which is comfortably below the sampling interval and thus compatible with real-time deployment. When averaging the estimated burst probability across hand movements (n = 41), we observed that beta burst probability was suppressed immediately prior to movement onset and remained low for nearly 1 s post-onset before rising again (Fig. 5). The narrow peak near 0 s appears to be a common occurrence in this dataset since in some subset of trials, a burst occurs in the immediate aftermath of movement initiation. The movement-related suppression and rebound is consistent with prior work on beta burst modulation around movement [12], supporting the validity of MSSR for detecting beta bursts in real data and informing stimulation timing. However, more extensive validation on real neural data would strengthen the evidence for the practical usefulness of the MSSR.

**Fig. 5.**
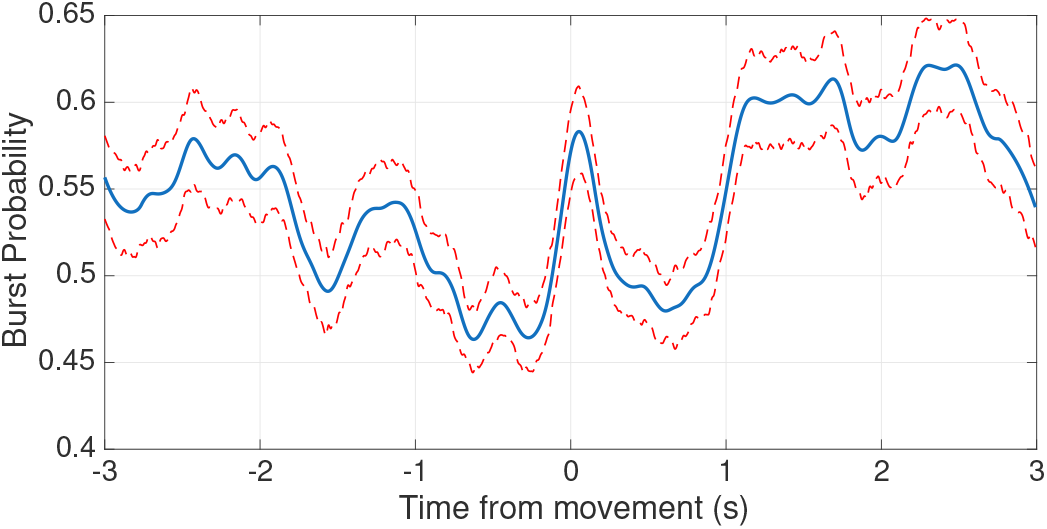
Average real-time estimated beta burst probability around hand movement onset. Red lines cover 95 % CI.

## IV. CONCLUSIONS

We introduced a novel state space method for real-time detection of oscillatory bursts, using Parkinsonian beta bursts as a clinically relevant application. The method provides a continuous, sample-by-sample estimate of burst probability and detects burst onset with low latency — on average 13 ms, which is faster than half a cycle of high-beta activity (17 ms at 30 Hz), and burst offset with near-zero latency (-12 ms) while contemporary methods took 100ms before they concluded a burst was over. For scale, in an aDBS setup, if burst detections were prolonged by 0.1 s per-burst and burst events occurred at 1 Hz, this would add 0.1 × 86,400 = 8,640 s, or 2.4 h, of stimulation time per day. In addition, burst detection was less sensitive to threshold selection than commonly used bandpass filtering-driven approaches, improving robustness for clinical deployment. Future work will extend the framework to simultaneous multi-band burst detection (e.g., concurrent beta and gamma bursts), as proposed in [13]. Overall, this approach offers a practical and accurate detection method for adaptive deep brain stimulation strategies that aim to time stimulation to pathological beta-burst events.

## Notes

### Competing Interest Statement

The authors have declared no competing interest.

## REFERENCES

[1] H. Bronte-Stewart, C. Barberini, M. M. Koop, B. C. Hill, J. M. Henderson, and B. Wingeier, “The stn beta-band profile in parkinson’s disease is stationary and shows prolonged attenuation after deep brain stimulation,” Experimental neurology, vol. 215, no. 1, pp. 20–28, 2009.

[2] Z. D. Langford and C. R. Wilson, “Simulations reveal that beta burst detection may inappropriately characterize the beta band,” Journal of Neurophysiology, 2025.

[3] G. Tinkhauser, A. Pogosyan, H. Tan, D. M. Herz, A. A. Kühn, and P. Brown, “Beta burst dynamics in parkinson’s disease off and on dopaminergic medication,” Brain, vol. 140, no. 11, pp. 2968–2981, 2017.

[4] G. Tinkhauser, A. Pogosyan, S. Little, M. Beudel, D. M. Herz, H. Tan, and P. Brown, “The modulatory effect of adaptive deep brain stimulation on beta bursts in parkinson’s disease,” Brain, vol. 140, no. 4, pp. 1053–1067, 2017.

[5] G. Karvat, A. Schneider, M. Alyahyay, F. Steenbergen, M. Tangermann, and I. Diester, “Real-time detection of neural oscillation bursts allows behaviourally relevant neurofeedback,” Communications biology, vol. 3, no. 1, p. 72, 2020.

[6] J. L. Busch, J. Kaplan, J. K. Behnke, V. S. Witzig, L. Drescher, J. G. Habets, and A. A. Kühn, “Chronic adaptive deep brain stimulation for parkinson’s disease: clinical outcomes and programming strategies,” npj Parkinson’s Disease, vol. 11, no. 1, p. 264, 2025.

[7] A. Wodeyar, M. Schatza, A. S. Widge, U. T. Eden, and M. A. Kramer, “A state space modeling approach to real-time phase estimation,” Elife, vol. 10, p. e68803, 2021.

[8] Y. Bar-Shalom, X. R. Li, T. Kirubarajan, et al., “Adaptive estimation and maneuvering targets,” Estimation with Applications to Tracking and Navigation: Theory, Algorithms and Software, pp. 421–491, 2001.

[9] A. Wodeyar, F. A. Marshall, C. J. Chu, U. T. Eden, and M. A. Kramer, “Different methods to estimate the phase of neural rhythms agree but only during times of low uncertainty,” Eneuro, vol. 10, no. 11, 2023.

[10] C. Zrenner, D. Galevska, J. O. Nieminen, D. Baur, M.-I. Stefanou, and U. Ziemann, “The shaky ground truth of real-time phase estimation,” Neuroimage, vol. 214, p. 116761, 2020.

[11] T. Merk, V. Peterson, W. J. Lipski, B. Blankertz, R. S. Turner, N. Li, A. Horn, R. M. Richardson, and W.-J. Neumann, “Electrocorticography is superior to subthalamic local field potentials for movement decoding in parkinson’s disease,” Elife, vol. 11, p. e75126, 2022.

[12] F. Torrecillos, S. He, A. A. Kühn, and H. Tan, “Average power and burst analysis revealed complementary information on drug-related changes of motor performance in parkinson’s disease,” npj Parkinson’s Disease, vol. 9, no. 1, p. 93, 2023.

[13] M. He, P. Das, G. Hotan, and P. L. Purdon, “Switching state-space modeling of neural signal dynamics,” PLoS Computational Biology, vol. 19, no. 8, p. e1011395, 2023.

